# Benchmarking Geometric Morphometric Methods: A Performance Evaluation for Gastropod Shell Shape Analyses

**DOI:** 10.64898/2026.02.23.707480

**Authors:** David Carmelet–Rescan, Gabriella Malmqvist, Luisa Kumpitsch, Beatrice Sammarco, Le Qin Choo, Roger Butlin, Francesca Raffini

**Affiliations:** Department of Biology and Evolution of Marine Organisms, Stazione Zoologica Anton Dohrn, Naples, Italy; Department of Marine Sciences, University of Gothenburg, Tjärnö Marine Laboratory, SE 452 96 Strömstad, Sweden; Department of Life Sciences and Biotechnologies, University of Ferrara, Ferrara, Italy; Ecology and Evolutionary Biology, School of Biosciences, The University of Sheffield, Sheffield, UK; Simon F.S. Li Marine Laboratory, School of Life Sciences, The Chinese University of Hong Kong, Hong Kong SAR, China

**Keywords:** *Littorina saxatilis*, Shape analysis, morphological evolution, adaptive divergence, marine conservation, Shell shape

## Abstract

Understanding morphological variation is crucial for the study of speciation and for conservation as it helps in assessing biodiversity and predicting responses to environmental changes. These approaches are broadly applicable but are especially valuable in marine environments, where species are often elusive, difficult to study, and face heightened threats from rapid environmental shifts. The marine snail *Littorina saxatilis* is notable for its extensive polymorphism in shell shape, size, and colour, with ecotypes that evolve in response to environmental forces including wave exposure and crab predation. Morphometric tools have been central to investigating the mechanisms driving this phenotypic divergence; yet, a direct comparison of their methodological efficacy is lacking. Here, we took advantage of *L. saxatilis’* ecotypes to contrast three morphometric approaches: elliptical Fourier analysis (EFA), landmarks-based geometric morphometrics (GM), and the growth-based model implemented in the *ShellShaper* software (SS). We assessed their clustering power, biological interpretability, robustness to measurement error and transferability among datasets. Our findings provide insights to guide method selection in studies aimed at exploring morphological variation: EFA is better suited for high-throughput screening and describing intermediate shapes; SS offers superior clustering power with highly interpretable growth parameters; and GM is best for detailed anatomical studies but is less efficient for large datasets. We provide guidelines to align method selection with specific research goals, balancing analytical efficiency with the required morphological and biological insight. By following this framework, researchers can ensure that robust morphological analysis is achieved, which is essential not only for elucidating mechanisms of adaptation and speciation but also for effective management and conservation of marine biodiversity.

## Introduction

The study of morphology, particularly the shape and form of organisms, is fundamental to understanding their biological and ecological roles. Shape and form can influence an organism’s function, its interactions with the environment, and its evolutionary trajectory. This foundational relationship makes morphometrics a powerful tool across multiple scales of biological inquiry. At the applied level, integrating morphometric data with genomic and ecological models enables researchers to diagnose vulnerabilities to climate change and track population health through morphological shifts over time (Christmas et al., 2022; Westbury et al., 2023) and identify adaptive traits that enhance survival under environmental stress (Galindo & Grahame, 2014). Morphometric data can serve as indicators of environmental stress and predictors of community responses to global change (Yan et al., 2024). Scaling up, large morphometric datasets help quantify morphological diversity and relate it to macroecological and macroevolutionary patterns, illustrating how environmental changes influence ecological roles (Serra et al., 2023). From an evolutionary perspective, morphology aids in predicting adaptive responses by distinguishing between specialist and generalist trajectories and evaluating the reliability of morphological adaptations to varying environmental conditions (Pilakouta et al., 2023; Renaud et al., 2005).

Among morphometric approaches, geometric morphometrics offers particularly critical insights into evolutionary processes—such as adaptation, divergence, and convergence—by providing tools for analyzing organismal shape and form with high precision (D. Adams et al., 2013; D. C. Adams & Felice, 2014; Rohlf, 1998). Consequently, geometric morphometric approaches have become the primary tool for quantifying and analyzing morphological variation in evolutionary biology, ecology, and conservation research, with applications ranging from detecting reproductive status in killer whales (Robinson & Visona-Kelly, 2025) to assessing seagrass stability through lucinid bivalve shells (Anderson et al., 2022) and showing that adaptive traits can be associated with polymorphic inversions in rough periwinkles (Koch et al., 2022). Crucially, this approach has proven particularly impactful in studying phenotypic variation and its correlation with genetic and environmental factors, thereby advancing our understanding of evolutionary dynamics and species’ resilience (Laffont et al., 2011; Mitteroecker & Gunz, 2009). Given the critical state of marine ecosystems, which are particularly vulnerable to climate change and human activities, the integration of morphometric, genomic, and ecological approaches is essential for identifying conservation priorities, studying adaptive processes, and protecting marine biodiversity (Lishchenko & Jones, 2021; Ramírez-Portilla et al., 2022).

Molluscs, particularly gastropods, exhibit extraordinary diversity in shell morphology, reflecting adaptations to varied environmental and sexual pressures, genetic drift, and developmental constraints (Johnson et al., 2019; Kocot et al., 2016). Within this group, the marine snail genus *Littorina* displays remarkable morphological and ecological diversity, making it an ideal model for investigating the genetic and environmental drivers of phenotypic evolution (K. Johannesson, 2016; K. Johannesson et al., 2024; Kess & Boulding, 2019). Among *Littorina* species, the rough periwinkle, *Littorina saxatilis*, is a prominent example due to its adaptability and distinct ecotypes that occupy different micro-habitats. A well characterised polymorphism in this species includes repeated parallel shell morphological variation driven by environmental pressures: the "Crab" snails, adapted to crab predation, are larger with thicker shells, whereas the "Wave" snails, adapted to high-energy shores, are smaller and thinner-shelled with a wider aperture. These ecotypes can be found close together on the same shore and in narrow contact zones, hybrids with intermediate phenotypes are observed (Butlin et al., 2014; K. Johannesson et al., 2010; Koch et al., 2022; Rolán-Alvarez, 2007). Consequently, *L. saxatilis* has become a premier model for studying adaptive divergence and ecological speciation (Garcia Castillo et al., 2024; K. Johannesson, 2016; Le Pennec et al., 2017).

Research on this system has demonstrated how divergent ecological pressures can drive phenotypic divergence and reproductive isolation (Butlin et al., 2014; Galindo & Grahame, 2014; Hollander et al., 2006; K. Johannesson et al., 2024). Studies of parallel phenotypic divergence across populations further investigate whether similar genetic mechanisms, including chromosomal inversions, underlie repeated adaptations, offering insights into convergent evolution (Galindo et al., 2009; K. Johannesson, 2016; Koch et al., 2022; Le Moan et al., 2024; Morales et al., 2019; Ravinet et al., 2016). Investigating this remarkable diversity in *L. saxatilis* has been profoundly shaped by advances in morphometric analysis. Early studies relied on linear measurements (e.g., shell length and width) to establish fundamental form-function relationships, such as the link between aperture size and crab predation pressure (B. Johannesson, 1986). The adoption of landmark-based geometric morphometrics approaches then enabled a more nuanced quantification of shell shape, facilitating more detailed comparisons among populations and ecotypes (Carvajal-Rodriguez et al., 2005; Conde-Padín et al., 2007). By analyzing multiple landmarks, this approach was pivotal in detecting subtle, ecologically relevant morphological variations driven by factors like wave exposure and predation (Grahame & Mill, 1989; Ravinet et al., 2016; Tirado et al., 2016). However, snail shells are arguably poorly suited to traditional geometric morphometric approaches. Lacking true anatomical landmarks and being intrinsically three-dimensional structures, most methods rely on 2D projections, pseudo-landmarks, and sliding landmarks that may not fully capture shell complexity. While the effectiveness of these methods is well established (Doyle et al., 2018), recent innovations have further expanded analytical possibilities. Notably, techniques like *ShellShaper* (Larsson et al., 2020) integrate landmarks and curves to model three-dimensional shell growth, capturing more complex geometries that are also more intuitive. This enhanced resolution has proven powerful for directly linking genetic variation to phenotypic outcomes in *L. saxatilis* (Koch et al., 2022).

Nevertheless, the reliability and biological insight gained from any morphometric study depend on controlling measurement error and making critical methodological choices. The selection of analytical techniques must balance statistical power with biological interpretability (Doyle et al., 2018), and stringent control of measurement error—whether random or systematic—is essential to avoid obscuring genuine biological signals (Collyer & Adams, 2024; Fruciano, 2016). It can be difficult to make a choice among these methods, as each offers distinct advantages for quantifying shell morphology, yet varies in complexity, sensitivity to error, and biological interpretability. Here, we compare three morphometrics methods that are all well-established but differ fundamentally in their approach to quantifying form:

1. **Elliptical Fourier Analysis (EFA):** This method captures the shell’s outline, using harmonic analysis (Fourier transformations) to model shapes, an approach that is particularly powerful in forms that lack discrete homologous landmarks (Dytham et al., 1992; Haines & Crampton, 2000);
2. **Geometric Morphometrics (GM):** Based on homologous landmarks and semi-landmarks, this approach uses Procrustes superimposition to isolate and compare shape variation after accounting for differences in size, position, and orientation (Gefaell et al., 2020; Mitteroecker & Gunz, 2009; Ravinet et al., 2016; Westram et al., 2018);
3. **The Growth-Based Model *ShellShaper* (SS):** This biologically informed approach uses a mathematical model of spiral shell growth, defined by a few key parameters, to compare forms. It represents a broader class of generative models applied to diverse morphological structures (Contreras-Figueroa & Aragón, 2023; Evans et al., 2021; Larsson et al., 2020; Mosleh et al., 2023).

By contrasting the efficiency, reliability, and discriminatory power of these techniques, this study aims to provide clear guidance for selecting morphometric methods in evolutionary, ecological and conservation studies of complex forms, especially in gastropods. We seek to identify which approach best captures the morphological diversity and ecological relevance of *L. saxatilis* ecotypes, thereby offering a methodological framework that can be applied to other systems where quantifying intricate shape is key to understanding the interplay between genetics, adaptation, and phenotype.

## Materials and Methods

### Sample collection

*L. saxatilis* specimens were collected on the island of Saltö, on the site of Ängklåvbukten (Sweden, 58°52’11.4"N 11°07’11.7"E) in spring 2024, from a ∼150 m Crab-Wave transect including the contact zone (Westram et al., 2018). A total of 30 snails were analysed, comprising ten individuals each of the Crab, Hybrid and Wave ecotypes. By analysing these samples, we can classify individuals according to transect position on the basis of their shell morphology, which is adapted to their local micro-environment.

### Study design

The robustness and repeatability of the methods were tested by generating replicate analyses for each individual to quantify variability from three sources: photography (two cameras, two pictures per individual), digitization event (two sessions), and digitization operator (two researchers) (Figure 1).

**Figure 1.**
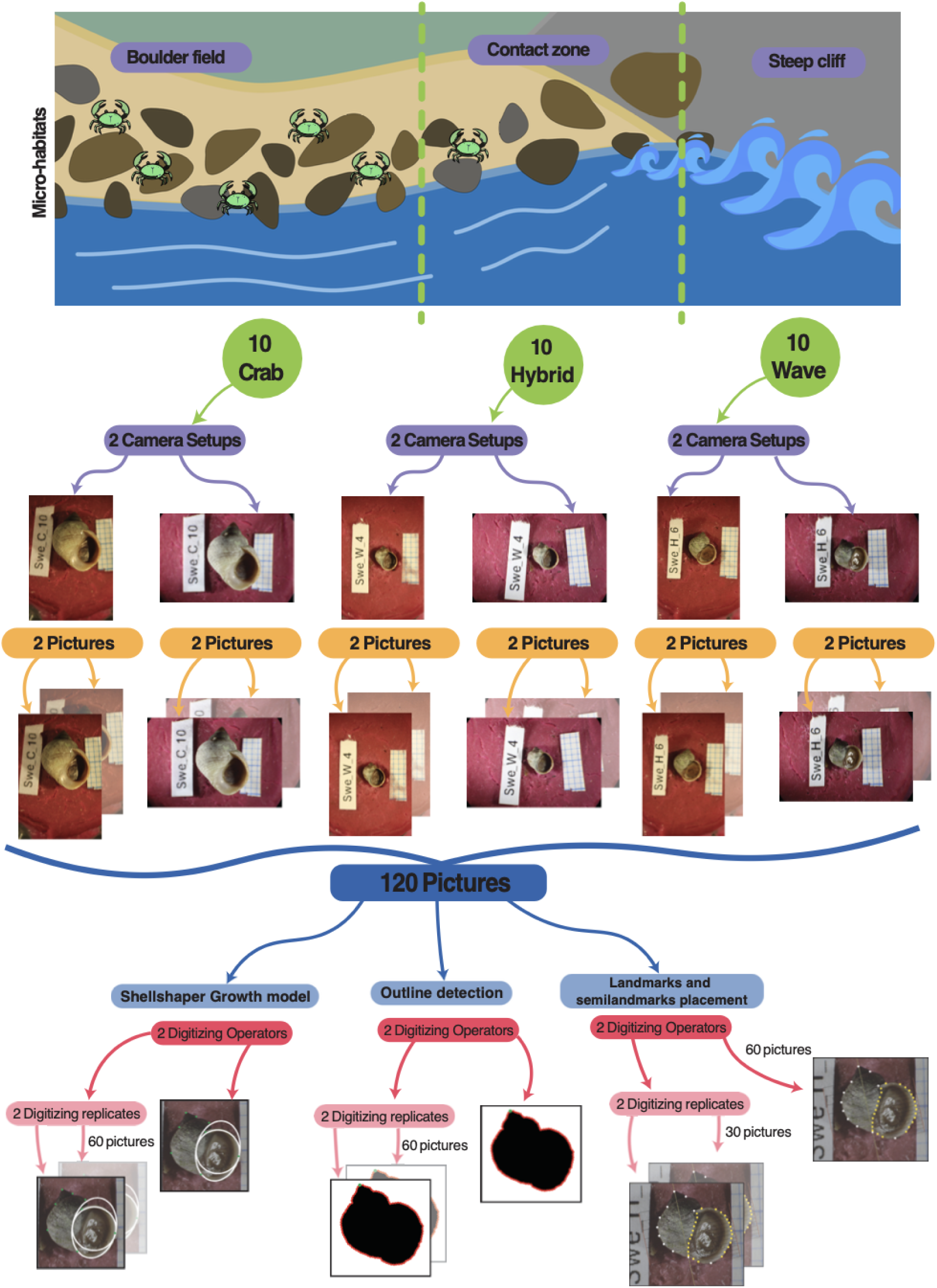
Experimental workflow and digitization schemes. The study design includes technical and digitization replicates for each morphometric method. The bottom row shows example images for the three digitization protocols: **SS**, highlighting the placement of landmarks and the defined aperture shape; **GM**, with the comb placement and the 38 landmarks capturing shell morphology; and **EFA**, featuring a shell outline with 1000 sampled points and the manually defined apex landmark.

Photographs of each specimen were obtained following a standardized protocol optimized for this species (Ravinet et al., 2016; Figure 1). To isolate sources of bias, each snail was photographed in four separate sessions using two different camera setups, with non-consecutive replicates for each setup. This design allowed us to distinguish variability arising from the imaging hardware *versus* specimen handling and placement. Two camera setups were used: Camera 1 consisted of a Canon EOS 5D Mark III mounted on an Olympus SZX16 stereoscope with auxiliary lens SDF PLAPO 0.5XPF, using an Olympus U-TV1X-2 adapter and Olympus U-CMAD3. Camera 2 consisted of a Canon EOS 600D mounted on an Olympus SZX12 stereoscope with auxiliary lens DF PLAPO 1XPF, using an Olympus U-TV0.5XC adapter.

### Digitization

SS digitization was performed by placing eight landmarks and defining the aperture shape with a circle and an ellipse, following the established software guidelines (Larsson et al., 2020). The software is a Matlab program and is accessible on this github repository: github.com/jslarsson/ShellShaper/tree/master. The two operators performing the digitization had differing levels of experience with the program (Figure 1).

For GS, we digitized 38 specific landmarks and semilandmarks along two comb templates placed over the shell aperture using the following configuration on Supplementary Figure 1.

This landmark configuration was adapted from established protocols, building upon a system developed for *L. saxatilis* (Ravinet et al., 2016) and a comb-based template used in other gastropod studies (Quenu et al., 2020). The comb branches intersections with the shell are defining the set of semi-landmarks, the template was applied in Adobe Photoshop v. 26.7 (Adobe Inc., 2019), and the 38 landmarks and semi-landmarks were digitized using the R package Stereomorph v. 1.6.7 (Olsen & Westneat, 2015). To capture the shell’s curvature accurately, 31 sliding semi-landmarks were placed between fixed landmarks. This step is critical for ensuring that the landmark set comprehensively represents the morphological features of interest.

In order to digitize the outline for the EFA, shells were isolated from the background using the Subject Detection tool in Adobe Photoshop, with manual refinements applied as needed. This process created a binary image, i.e., a black shell outline on a white background. Outlines were then imported into R (4.5.1) using the *Momocs* package v. 1.4.1 (Bonhomme et al., 2014), smoothed, and uniformly sampled with 1000 points. Finally, the apex was manually defined for each shell to standardize orientation in subsequent analyses (Figure 1).

The digitization protocol incorporated replication to quantify two sources of error: inter-operator variability and intra-operator repeatability. For SS and EFA, the complete set of 120 pictures was digitized independently by two operators. Furthermore, 60 pictures were re-digitized by one operator to isolate measurement repeatability. For GM, only 60 pictures were digitized by two operators, and a smaller subset of 30 pictures was digitized twice by one operator for the same purpose (Figure 1).

Additional guidelines to the digitization protocols mentioned in this article can be found in this more detailed protocol: https://www.protocols.io/private/CDEDFDB09DE711F0AA170A58A9FEAC02

### Processing

Elliptical Fourrier Analysis was performed using the *Momocs* package (Bonhomme et al., 2014). Shell outlines were extracted from processed images using the import_jpg function, which also performed outline smoothing and uniformly sampled 1000 points per outline. The apex was then manually defined for each shell to serve as a homologous point for alignment. Outlines were aligned using a Procrustes superimposition, a standard geometric morphometric method that removes variation in position, scale, and rotation (Gower, 1975). In this implementation, *Momocs* aligned the start point of outlines by sliding them to the defined apex landmark, ensuring homologous orientation for subsequent comparisons. Following alignment, we performed an Elliptical Fourier Analysis (EFA) on the outline coordinate matrix, decomposing each shape into a series of 50 harmonics to capture detailed morphological variation (Ferson et al., 1985). After removing outliers using the *which_out* function of *Momocs*, the resulting Fourier coefficients were subjected to a Principal Component Analysis (PCA) to reduce dimensionality. Principal components that cumulatively accounted for more than 95% of the total variance were retained for all downstream analyses (Figure 2).

**Figure 2.**
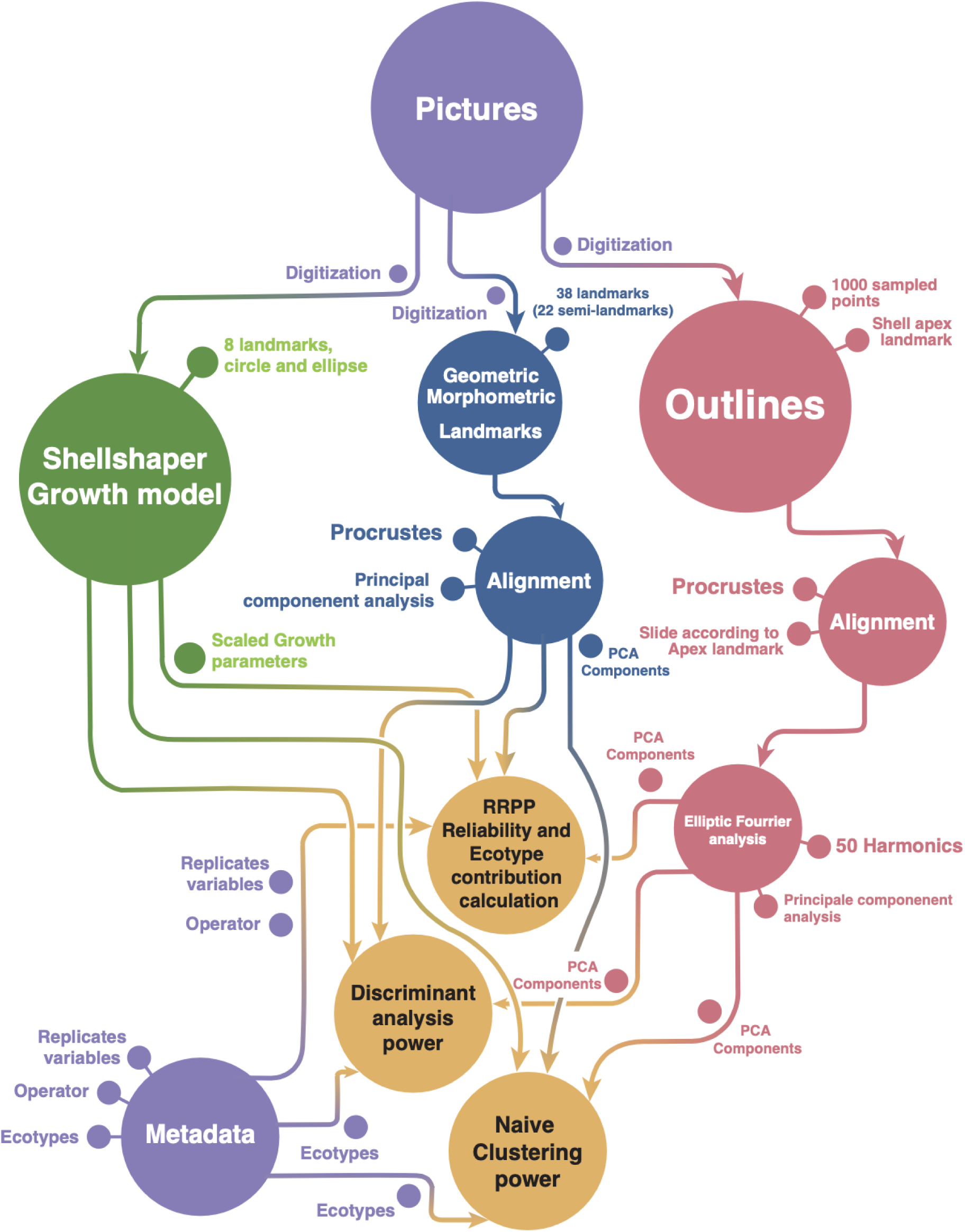
Data analysis workflow. Flowchart of the sequential steps from raw digitized data through performance analyses for the three morphometric methods. The big circles are the main steps and small circles on top of the arrows correspond to the type of data that is generated or used for each analysis.

For SS, the *Shellshaper* software directly outputs seven growth parameters (gw: growth width; gh: growth height; r0: radial distance from the origin; h0: vertical distance from the origin; a0: circle radius; c0: circlipse extreme point; apAngle: angular parameter, see Larsson et al., 2020 for full descriptions). Outliers were identified and removed using a 99% cutoff on mahalanobis distance from center. This data set was then imported and used directly in subsequent reproducibility and clustering analyses.

The GM landmark coordinates obtained from Stereomorph digitization were processed within the R package *geomorph* v. 4.0.10 (D. Adams et al., 2017). The data were first subjected to a Generalized Procrustes Analysis (GPA) to remove variation due to position, scale, and orientation and isolate true shape variation. After identifying and removing outliers, we performed a Principal Component Analysis (PCA) on the Procrustes coordinates via the geomorph::gm.prcomp function. The PCs representing cumulatively more than 95% of variance were then used in downstream statistical evaluations, including assessments of reproducibility and ecotype clustering.

### Performance assessment

Each morphometric technique was assessed across five metrics: processing efficiency (digitization time), reliability (importance and source of errors), biological signal strength, discriminant power, and naive clustering success (Table 1).

**Table 1.** Performance metrics used to evaluate the techniques in the study.

| Performance metric | Description | Input data | Output metric |
| --- | --- | --- | --- |
| Processing efficiency | Measures the average processing time for each samples | Time recorded for different stages of each technique | Processing time per sample |
| Reliability | Quantifies the reliability of results across technical replicates | Technical replicates (photos, cameras, operators, digitization events) | Intraclass Correlation Coefficient (ICC) |
| Biological signal strength | Quantifies the phenotypic distinction between groups using a linear model | Shape data and biological grouping (Crab, Wave, Hybrid) | Group signal (ratio of sum of squares) |
| Discriminant power | Evaluates the ability of each method to classify samples accurately according to ecotype using linear discriminant analysis | Principal components or scaled shape parameters | Classification Error and Brier Score |
| Naive clustering success | Tests how well the morphometric techniques can directly recover ecotypes from morphology using clustering algorithm | Principal components or scaled shape parameters of Crab and Wave samples | Adjusted Rand Index (ARI) |

For the first metric, we measured the total processing time to complete digitization for each technique processed by the same operator with a similar experience level in all methods. For EFA, time was recorded for three stages: Photoshop processing, automated outline detection, and manual apex placement. SS timing covered a single session of 90 samples. GM time encompassed comb placement in Photoshop and landmark digitization in Stereomorph.

The goal of the second metric was to quantify the consistency of results across technical replicates (photos, cameras, operators, digitization events). While repeatability focuses on the stability of measurements, reliability—quantified here using the Intraclass Correlation Coefficient (ICC)—integrates measurement error with population variability. This provides a more comprehensive assessment of a method’s stability in comparative studies (Bartlett & Frost, 2008). We conducted this analysis using the RRPP v. 2.1.2 package in R (Collyer & Adams, 2018). This package uses residual randomization in a permutation procedure to generate empirical probability distributions for evaluating model effects. For our model, which included the biological group (Crab, Wave and Hybrids) as a grouping factor, we used the measurement.error function to compute measurement errors. The ICCstats function was then used to calculate the ICC (Armqvist & Martensson, 1998; Fruciano, 2016), which quantifies the proportion of total variance due to true subject differences *versus* measurement error. A higher ICC indicates greater reliability and lower measurement noise. A key advantage of the RRPP framework for ICC calculation is that it mitigates a known limitation of the classic ICC: its direct dependence on the population variance of the specific sample (Cardini, 2020; Collyer & Adams, 2024; Fox et al., 2020). By using a permutation procedure, RRPP provides a more robust and generalizable estimate of reliability that is less influenced by the particular group variances in the study sample. For each technical replicating factor the fraction on its associated sum of squares on the total sum of squares is computed to assess each effect individually.

In the third metric, the phenotypic distinction between groups (Crab, Wave, Hybrid) was quantified by fitting a linear model with RRPP::lm.rrpp, where shape was the response variable and biological grouping was the predictor. An ANOVA on this model provided sums of squares, from which we calculated the effect for groups as the ratio of the among-group sum of squares to the total sum of squares. This ratio, which we term the ’group signal’, represents the proportion of total variance attributable to differences between groups.

For the fourth metric, we performed a discriminant analysis to evaluate the ability of each morphometric technique to classify samples accurately into their known groups (Crab, Wave, Hybrid). This assesses how effectively each method distinguishes groups based on morphological differences. The analysis was conducted using the MclustDA function from the *mclust* R package (v6.1.1) (Fraley et al., 2017). This method fits a discriminant analysis model based on Gaussian finite mixture modeling. We specified the "EDDA" (Eigenvalue Decomposition Discriminant Analysis) model, which is well-suited for high-dimensional data like principal components or scaled shape parameters. The main ecotypes and hybrids were used as the grouping variable. To assess the model’s robustness, we performed 10-fold cross-validation using the cvMclustDA function. Model performance was evaluated using two key metrics derived from the cross-validation: i) Classification Error: The proportion of misclassified samples; and ii) Brier Score: A measure of the accuracy of the probabilistic predictions (Brier, 1950). Lower values for both metrics indicate a more effective biological group discrimination.

The fifth metric involved a naive clustering analysis, i.e., grouping samples into distinct clusters based on their morphological characteristics without prior knowledge of their ecotypes. This analysis tests how well the morphometric techniques can naturally recover ecotypes. Clustering was based on the PCs (or scaled shape parameters for SS) that cumulatively accounted for more than 95% of the variance. We used the mclust function for model-based clustering, which fits a Gaussian finite mixture model. To optimize the model, we employed the clustvarsel function (clustvarsel v2.3.5; (Scrucca & Raftery, 2018) for variable selection, constraining the model to retain the first principal component. The analysis was specified to fit a two-cluster solution, corresponding to the two distinct ecotypes (Crab and Wave). Hybrid snails were excluded from this analysis because their intermediate morphology makes them difficult to classify reliably using naive clustering methods.The agreement between the resulting clusters and the known ecotype labels was quantified using the Adjusted Rand Index (ARI, Steinley, 2004). The ARI measures the similarity between two clusterings, with a value of 1 indicating perfect agreement and 0 indicating random assignment. A higher ARI signifies a greater ability of the technique to recover the known ecotype.

### Correlation analysis

To examine how EFA compares to SS growth parameters, we performed a correlation analysis between PCs derived from EFA and SS. Due to deviation from normality, Spearman correlation coefficients were computed between each of the PCs from the EFA that cumulatively accounted for more than 95% of the variance and the SS growth parameters, p-value were adjusted using Bonferroni correction. This analysis allowed us to quantify the strength and direction of the linear relationships between overall EFA output and specific growth traits.

All the analyses in this study were performed on a MacBook Pro laptop with M3 Pro chip, 18 GB RAM.

## Results

### Replicates

To quantify technical variance, we processed 30 individuals through a replicated workflow assessing photographic, digitization, and operator effects (Figures 1 and 2). Sample sizes for each morphometric technique (EFA, SS, GM) and replication step are specified in Table 2.

**Table 2:** Spatial and temporal scope of the study and definition of observational units.

| Scale of interference | Experimental unit | Number of replicates | Response variables |
| --- | --- | --- | --- |
| Three biological groups (Crab, Hybrid, Wave ecotypes) | Individual *L. saxatilis* | 10 specimens per biological group | Shell Shape |
| Three Morphometric Methods | Digital representation of a snail shell shape | 30 pictures per technique | Processing Time |
|  |  | 2 camera setups per sample | Reliability |
|  |  | 2 pictures per camera setup | Biological Grouping Signal |
|  |  | 2 digitizing operators per technique | Group Discrimination |
|  |  | 2 digitizing replicates for 30-60 pictures | Method Correlation |

### Processing Time

We evaluated the performance of each technique by measuring the digitization processing time per sample. EFA was the fastest method (12.8 seconds per samples on average), followed by SS (102.7 seconds per samples on average) and GM (202.6 seconds per samples on average), which required significantly more time. It should be noted that processing times can be influenced by computational resource and operator experience.

### Reliability

The Intraclass Correlation Coefficient (ICC), which quantifies measurement consistency across replicates, was computed for each technique, values closer to 1 indicate high repeatability, the score corresponds to the fraction of variation among groups over the total phenotypic variance of the population (Stoffel et al., 2017). EFA demonstrated excellent reliability (ICC = 0.927), followed by SS (ICC = 0.863) and GM(ICC = 0.717). This indicates that the first two methods produce highly consistent results, whereas the last one exhibits greater variability between replicates of all types.

Analysis of variance components revealed that the primary difference in reliability among methods stemmed from error introduced during the digitization phase. This includes variability from both different operators and technical digitization replicates (Figure 3). EFA technique’s high reliability is likely attributable to its largely automated digitization process, which minimizes the potential for human-induced error. In all techniques the measurement errors represent less than 10% of the variance attributed to the difference between individuals showing decent reliability across techniques (EFA: 1.04%, SS: 6.20%, GM: 7.05%).

**Figure 3.**
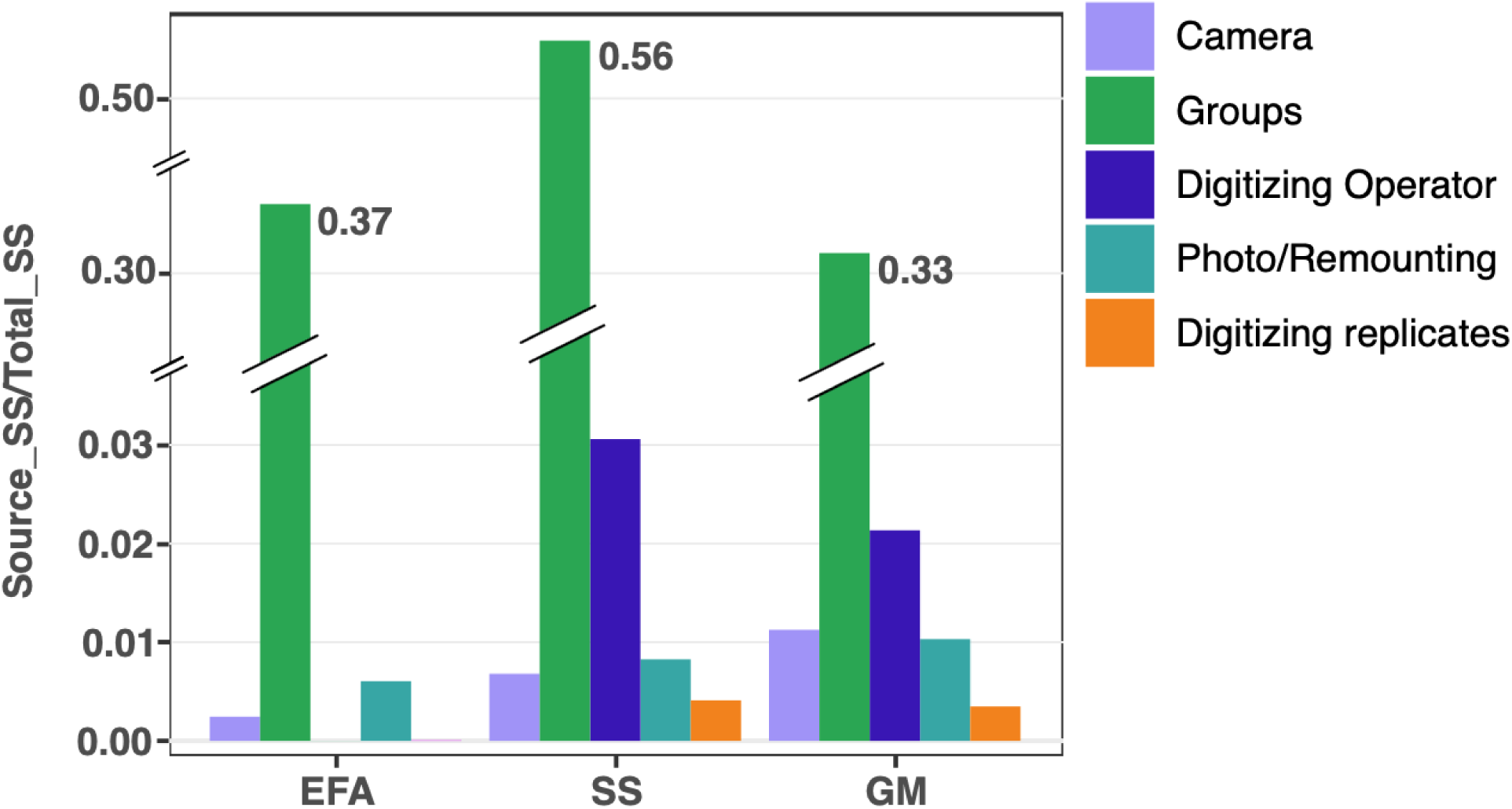
Relative impact of technical error sources on morphometric data. Bars show the proportion of total variance (sum of squares from RRPP) attributed to different stages of data acquisition and groups for the Elliptic Fourier Analysis (EFA), Shellshaper (SS), and Geometric Morphometric (GM) techniques. Digitizing operator and Digitizing replicates values of EFA are below 1e-4 though not showing on the graph.

**Figure 4.**
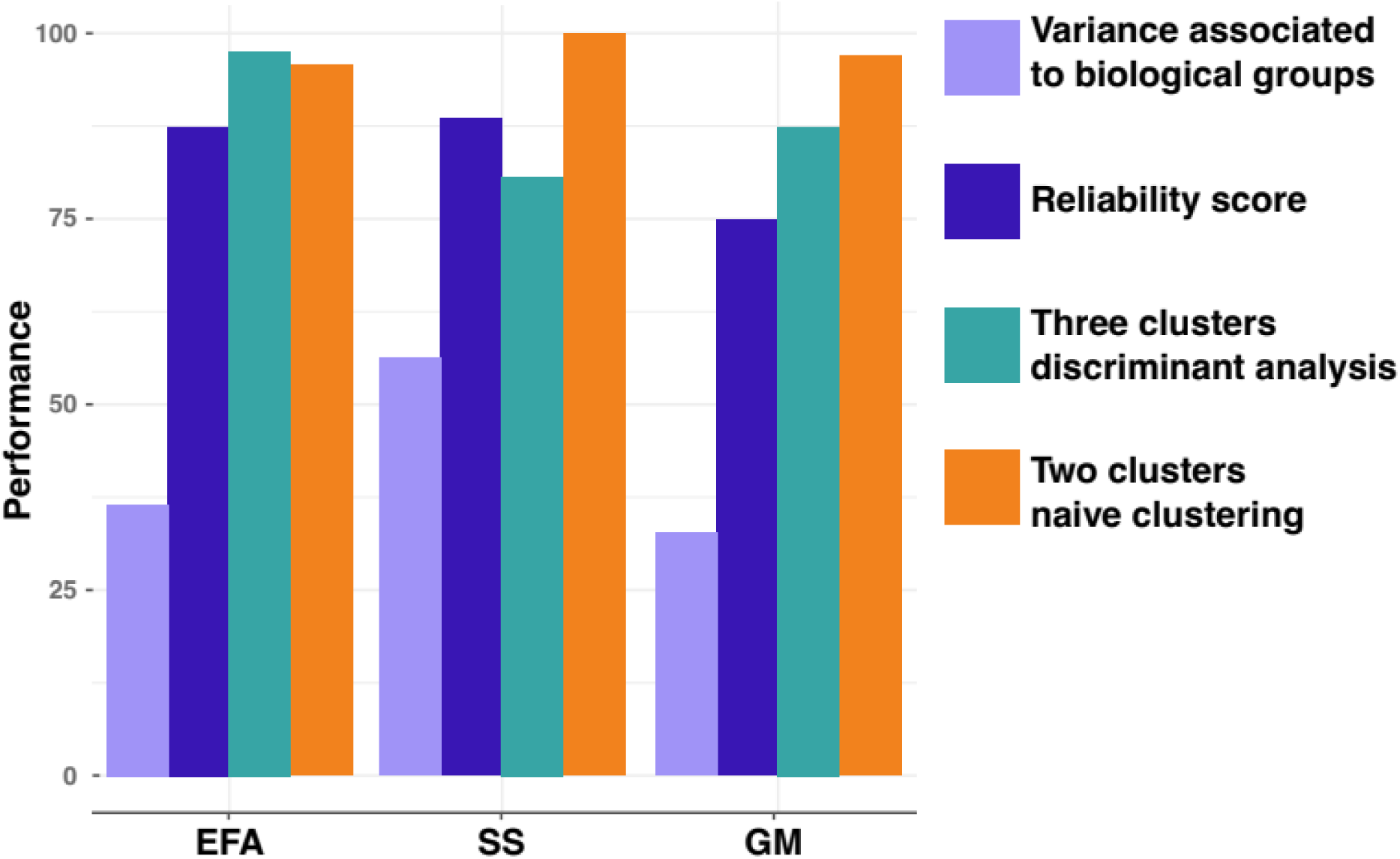
A multi-criteria assessment of morphometric technique performance, variance associated to biological groups (quantifying ecotype discrimination), Reliability score (ICC), Three clusters discriminant analysis (classification error), and two clusters naive clustering agreement (adjusted rand index).

### Biological grouping signal

The biological groups signal—the percentage of morphological variance attributed to morphological differences among the three groups—revealed that SS (56.2%) carried a stronger signal than both AFA (36.5%) and GS (32.6%) techniques.

### Group Discrimination

The effectiveness of each technique in discriminating between “Crab”, “Wave” and “Hybrids” was evaluated using discriminant analysis and naive clustering.

The classification error in the discriminant analysis was lowest for EFA (2.4%), followed by GS (7.5%) and SS (18.3%). This pattern was mirrored in the Brier scores (EFA: 0.018, GS: 0.065, SS: 0.140), confirming that EFA is the most effective for supervised classification into the known Crab, Wave, and Hybrid groups (Supplementary Figure 2).

The agreement between unsupervized clusters and true ecotypes, measured by the Adjusted Rand Index (ARI), was highest for SS (100%), followed by GM (97%) and EFA (95.7%). This indicates that SS is the most effective technique for recovering the natural morphological grouping between the Crab and Wave ecotypes without prior group information.

### Method Correlation

Significant correlations were observed between the PCs from EFA and SS (Fig. 5). EFA PC1 was strongly and significantly correlated with all SS growth variables, reflecting the fundamental property of PCA in which the first principal components capture the largest portions of variance of the Crab-Wave differences. The other EFA PCs exhibited more variable relationships, showing a complex interplay between overall shell shape and specific growth processes.

**Figure 5:**
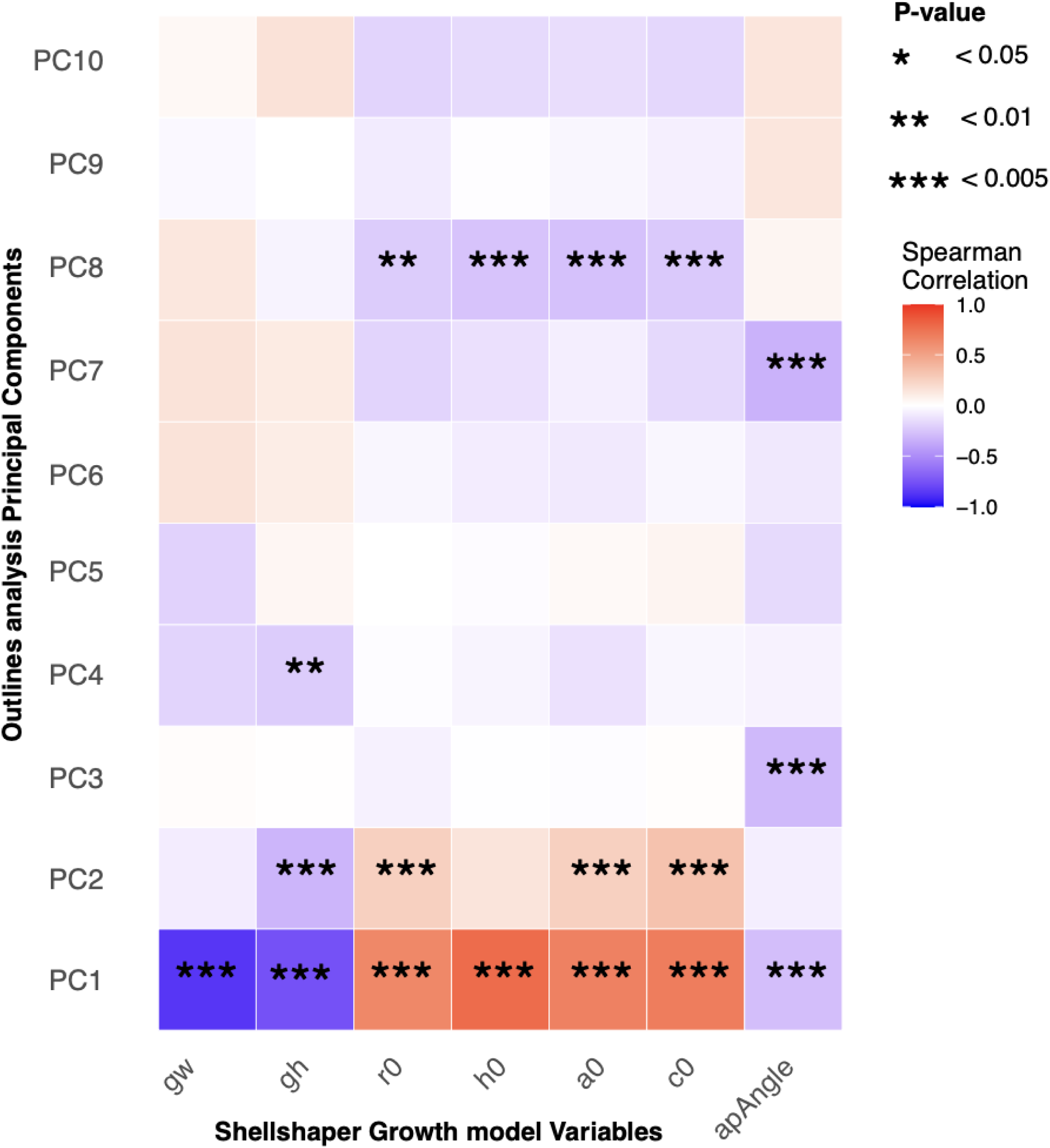
Correlations between EFA Principal Components and SS growth parameters. Color intensity indicates the strength of the correlation, while asterisks show statistical significance.

## Discussion

Phenotypes such as shape are the primary targets of natural selection, making their precise measurement fundamental to evolutionary ecology and conservation, particularly relevant to marine ecosystems, where species are often elusive, difficult to study, and face heightened threats from rapid environmental shifts. Different research questions may require different morphometric techniques. For instance, exploring general differences between ecotypes may benefit from a method that captures broad morphological variation, whereas identifying key distinguishing features may require a more focused approach. To determine which approach best suits different research objectives, we benchmarked three distinct methods — Elliptic Fourier Analysis (EFA), *ShellShaper* (SS), and Geometric Morphometrics (GM) — using the exuberant shell shape variation observed in *L. saxatilis*. Our results provide a practical framework to help researchers align their methodological choice with core research priorities, whether they emphasize analytical speed, biological interpretability, or the resolution of fine-scale morphological detail.

### Technique-dependent trade-offs

Within the Swedish Crab-Hybrid-Wave system of *L. saxatilis*, EFA emerged as the most efficient and reliable high-throughput tool. It achieved the lowest classification error (2.4%) and demonstrated high consistency across replicates (ICC = 0.927; Fig. 5), making it well-suited for batch-processing large datasets. Hybrids show substantial morphological variation, and not all are intermediate between parental ecotypes. This variation makes clustering hybrids particularly challenging, as some hybrids may be morphologically very similar to the true ecotypes. The choice of method depends on the specific research question. For example, if the goal is to explore general differences between ecotypes, a method that captures broad morphological variation, such as EFA, may be more suitable. Conversely, if the goal is to identify key distinguishing features, a more focused approach such as SS may be necessary. The EFA "naive" sampling of the entire shell outline provides a broad feature set, enabling discriminant analysis to capture subtle and complex morphological variations beyond the primary axis of ecotype divergence. A primary limitation, however, is its abstract output, which depends on PCA for interpretation. Furthermore, the method lacks flexibility; integrating new data requires restarting the entire pipeline from initial images or contours, as the PCA issued from Elliptical Fourier transformation analysis are defined by the variance of the original training dataset.

In contrast, SS excels as a targeted, hypothesis-driven tool for ecotype discrimination, resulting in optimal clustering (100% Adjusted Rand Index) and capturing the highest proportion of the group signal (56.2%). Additionally, the occasional association of hybrids with parental ecotypes in the discriminant analysis is not necessarily problematic, as some hybrids are actually morphologically very similar to the pure ecotypes. Its primary advantage lies in generating standardized, absolute growth parameters rather than relative scores from PCA. This provides direct biological interpretability and facilitates cross-study comparability without the need of repeating the entire data collection steps (Larsson et al., 2020). However, this focus comes at a cost: its performance in hybrid identification was inferior, and it depends on licensed software. Importantly, its landmark system is not universally applicable; it works well for similarly shaped molluscs (Jackson et al., 2021) but could be unsuitable for other species. This is the case of the congeneric flat snails *L. fabalis* and *L. obtusata,* since their shell geometry prevents simultaneous high-resolution visualization of both the aperture and the apex in a single two-dimensional image. Additionally, SS offers high reproducibility when operated by a trained user but requires specific expertise to master the precise placement of landmarks and curves. In particular, defining the aperture using the combination of a circle and an ellipse demands considerable practice, as suboptimal positioning can significantly bias the results. To ensure consistency and user proficiency, in addition to the official software manual, we recommend consulting the detailed protocol accompanying this study (https://www.protocols.io/private/CDEDFDB09DE711F0AA170A58A9FEAC02). Furthermore, practicing with our benchmark dataset (available at: https://figshare.com/projects/_b_Benchmarking_Geometric_Morphometric_Methods_A_Performance_Evaluation_for_Gastropod_Shell_Shape_Analyses_b_/265810) is strongly advised to validate correct landmark placement before applying the method to new data.

The strong correlation between SS growth parameters and EFA PC1 confirms that both methods capture the primary axis of shell shape variation. However, the subsequent EFA PCs captured orthogonal, additional information. It was this additional shape variation—unmeasured by the targeted SS protocol—that was leveraged to identify hybrids, demonstrating the advantage of a high-dimensional approach for comprehensive shape capture. It is important to add that the capacity of EFA to separate hybrids might be less robust biologically than the SS growth parameters placement of hybrids as simply intermediate with some overlap with parental ecotype distributions.

GM provided high anatomical detail but proved to be the least practical for large-scale studies. It exhibited the highest susceptibility to operator error and significant inter-observer variability, which directly undermined its reliability and analytical efficiency. Furthermore, like EFA, GM relies on PCA for data reduction, resulting in similarly abstract shape variables. These methodological constraints make cross-study comparisons tricky without proper estimation of operator bias or complete re-digitization of all specimens, severely limiting the method’s reproducibility and scalability.

Due to a focus on other effects, and design limitations, this study did not assess the effect of the operator on mounting the snail and taking the picture, which could introduce variability in the data collection process. This should be considered in future work.

### Choosing a Morphometric Technique: A Performance-Based Guide

Our benchmarking of SS, EFA, and GM clarifies which approach best suits different research objectives (Table 3). For studies aiming to discriminate defined morphotypes such as our Crab and Wave snails with directly interpretable growth parameters, SS is the optimal tool, despite its requirement for operator training. When the objective is the high-throughput analysis of complex morphological variation—such as analyzing intermediate forms like our Hybrid snails—EFA is superior due to its high reliability, efficiency, and rich, high-dimensional data output. The anatomical precision of GM is offset by its high susceptibility to operator error and consequent impracticality for large-scale studies. Consequently, no single technique is universally optimal; the choice should instead be guided by whether the research priority is targeted discrimination, high-throughput versatility, or anatomical precision.

**Table 3:** Comparison of three morphometric techniques analysed in this study.

| Criteria | ShellShaper | Elliptical Fourier Analysis | Geometric Morphometrics |
| --- | --- | --- | --- |
| Reliability | Requires trained operators. | Less prone to operator errors, dependent on photo quality | Lowest, influenced by the number of landmarks, require trained operators. |
| Data Comparability | Growth parameters are directly comparable across studies, prone to strong operator effect. | Requires outline coordinates and re-analyzing for comparison with other datasets. | Comparisons can be done from similar landmark data sets, but prone to strong operator effect. |
| Time Efficiency | Moderately efficient with a trained operator | Very fast and suitable for batch processing | Slow but may be improved with AI_based landmark placement. |
| Interpretability | Growth parameters directly linked to shell shape, intuitive and easy to integrate | Requires dimension reduction process. | Requires dimension reduction process. |
| Ease of Use | Moderate; requires specific training | High; minimal training required. | Moderate; requires specific training |
| Morphological fidelity | High for targeted morphological signals, biologically relevant but ignoring all details outside the model. | Moderate; captures precise outline but forgo any detail inside the shape (aperture). | High; captures precisely any morphological feature that was targeted by the landmark system. |
| Error Sources | Operator variability, positioning quality | Photo quality, positioning quality | Operator variability, positioning quality |
| Computing Dependency | Low computing power requirement but specialized and licensed software required (Matlab). | Requires good image processing software, processing speed can be dependent on computing power. | Low requirements on computing power and many available software options. |
| Practicality for Large Datasets | Moderately efficient with trained operator | Very efficient for large datasets using image processing automations. Dependent on computing power. | Slow processing speed is very dependent on the number of landmarks, path toward machine learning landmark placement. |

These findings illustrate a broader geometric morphometrics methodological dichotomy in analytical paradigms: one that leverages high-throughput data and powerful statistical tools like dimensionality reduction (Boyer et al., 2015; Collyer et al., 2015; Goswami et al., 2019), and a more hypothesis-driven approach that prioritizes biologically informed models tailored to specific questions (Cardini, 2020; Larsson et al., 2020; Contreras-Figueroa & Aragón, 2023; Pascoletti, 2022). Such biologically-driven models (e.g., SS) prioritize interpretability by linking morphology directly to underlying generative processes like growth (Evans et al., 2021; Larsson et al., 2020). They avoid the "black box" of multivariate statistics and yield absolute measurements that promote cross-study standardization (Cardini, 2020). However, their targeted nature may miss morphological signals outside their predefined scope. In contrast, high-density sampling approaches (e.g., EFA, GM) offer a comprehensive, data-driven representation of form, enhancing the detection of subtle and unanticipated shape variation (Goswami et al., 2019). This descriptive power comes with interpretability challenges, increased susceptibility to measurement error, and a reliance on dimension-reduction techniques that can obscure biological signals (Cardini et al., 2019). Our study thus demonstrates a fundamental trade-off: targeted biological interpretability *versus* high-dimensional descriptive versatility, where the optimal method is dictated by the research question.

In conclusion, our benchmarking study shows that methodological choice critically governs the trade-off between analytical efficiency, statistical reliability, and biological insight. We propose a strategic selection framework for future morphometric studies: SS is optimal for testing specific hypotheses related to shell growth in shells with geometry similar to *Littorina saxatilis* and also to study specifically how different are pre-defined morphotypes; EFA excels in high-throughput discovery of complex morphologies and is the most agnostic approach; and GM is best suited for detailed anatomical studies where scale is not limiting.

The importance of this framework extends beyond methodological considerations, as morphological variation itself holds critical ecological and evolutionary significance. For example, selecting the appropriate method is particularly crucial given the role morphological data play in understanding organismal responses to environmental change. The practical implications of these methodological distinctions become especially relevant in the context of rapid environmental change. This framework for method selection takes on heightened importance when we consider that accurate morphological quantification underpins our ability to detect and interpret organismal responses to environmental stress. As morphological shifts serve as a sensitive record of adaptive and plastic responses to environmental pressures (Albano et al., 2025) and a vital early-warning indicator of stress (Clark et al., 2013; Mayk et al., 2022), the rigorous quantification of this variation becomes essential. Our findings, therefore, offer a blueprint for integrating diverse morphometric data—from the general applicability of EFA and GM to the specialized utility of SS in gastropods with similar shell geometries (Larsson et al., 2020)—into a cohesive eco-evolutionary framework. As global change accelerates, this strategic alignment of tools with research questions is paramount for linking genotype to phenotype, assessing adaptive capacity, and forecasting population resilience in a rapidly changing world.

## Acknowledgment

We are grateful to Jenny Larsson and Eva Koch for indications on the *ShellShaper* software, Anja Westram and Zuzanna Zagrodzka for material and advice on the picture process. ChatGPT (GPT-5.1) and Mistral AI (Mistral Large 2) were used in the preparation of this manuscript only to identify and correct grammatical errors and improve the clarity of the writing. DCR and BS were supported by Progetti di Rilevante Interesse Nazionale (PRIN) 2022 PNRR, funded by the Ministero dell’Università e della Ricerca and European Union - NextGenerationEU, Decreto Direttoriale n. 1409 del 14-9-2022, CUP C53D23007100001, Project Code: P202229JBC to FR.

## Data availability

Raw data is currently available here: https://figshare.com/projects/_b_Benchmarking_Geometric_Morphometric_Methods_A_Performance_Evaluation_for_Gastropod_Shell_Shape_Analyses_b_/265810

The custom codes and intermediate files are archived in the GitHub repository: https://github.com/dcarmelet/Mollusc_Morphometrics.

The digitization protocols are publicly available in protocols.io: https://www.protocols.io/private/CDEDFDB09DE711F0AA170A58A9FEAC02.

## Authors contributions

Conceptualization: DCR, FR, RB; Project supervision: FR; Data collection: DCR, GM, LK, BS; Analysis: DCR, FR, LCQ; Discussion: DCR, FR; Writing - original draft: DCR, FR; Writing - review and editing: FR, DCR, RB, GM, LK, LCQ; Funding: FR, GM.

## References

Adams, D. C., & Felice, R. N. (2014). Assessing Trait Covariation and Morphological Integration on Phylogenies Using Evolutionary Covariance Matrices. PLOS ONE, 9(4), e94335. 10.1371/journal.pone.0094335

Adams, D., Collyer, M., Kaliontzopoulou, A., & Sherratt, E. (2017). geomorph: Geometric Morphometric Analyses of 2D/3D Landmark Data (Version 3.0.5) [Computer software]. https://CRAN.R-project.org/package=geomorph

Adams, D., Rohlf, F. J., & Slice, D. (2013). A Field Comes of Age: Geometric Morphometrics in the 21st Century. Hystrix, 7–14. 10.4404/hystrix-24.1-6283

Adobe Inc. (2019). *Adobe photoshop* (Version CC 2019) [Computer software]. https://www.adobe.com/products/photoshop.html

Albano, P. G., Kallmeyer, S. K. C., Vetina, A. A., Tibiriçá, Y., de Abreu, D. C., & Modica, M. V. (2025). A unique historical baseline uncovers harvesting impacts on intertidal molluscs at Inhaca Island, southern Mozambique. ICES Journal of Marine Science, 82(11), fsaf206. 10.1093/icesjms/fsaf206

Anderson, L. C., Long-Fox, B. L., Paterson, A. T., & Engel, A. S. (2022). Live and Live-Dead Intraspecific Morphometric Comparisons as Proxies for Seagrass Stability in Conservation Paleobiology. Frontiers in Ecology and Evolution, 10. 10.3389/fevo.2022.933486

Armqvist, G., & Martensson, T. (1998). Measurement error in geometric morphometrics: Empirical strategies to assess and reduce its impact on measures of shape. Acta Zoologica Academiae Scientiarum Hungaricae, 44(1–2), 73–96.

Bartlett, J. W., & Frost, C. (2008). Reliability, repeatability and reproducibility: Analysis of measurement errors in continuous variables. Ultrasound in Obstetrics & Gynecology, 31(4), 466–475. 10.1002/uog.5256

Bonhomme, V., Picq, S., Gaucherel, C., & Claude, J. (2014). Momocs: Outline Analysis Using R. Journal of Statistical Software, 56, 1–24. 10.18637/jss.v056.i13

Brier, G. W. (1950). Verification of Forecasts Expressed in Terms of Probability. Monthly Weather Review, 78(1), 1–3. 10.1175/1520-0493(1950)078%253C0001:VOFEIT%253E2.0.CO;2

Butlin, R. K., Saura, M., Charrier, G., Jackson, B., André, C., Caballero, A., Coyne, J. A., Galindo, J., Grahame, J. W., Hollander, J., Kemppainen, P., Martínez-Fernández, M., Panova, M., Quesada, H., Johannesson, K., & Rolán-Alvarez, E. (2014). Parallel Evolution of Local Adaptation and Reproductive Isolation in the Face of Gene Flow. Evolution, 68(4), 935–949. 10.1111/evo.12329

Cardini, A. (2020). Less tautology, more biology? A comment on “high-density” morphometrics. Zoomorphology, 139(4), 513–529. 10.1007/s00435-020-00499-w

Carvajal-Rodriguez, A., Conde-Padín, P., & Rolan-Alvarez, E. (2005). DECOMPOSING SHELL FORM INTO SIZE AND SHAPE BY GEOMETRIC MORPHOMETRIC METHODS IN TWO SYMPATRIC ECOTYPES OF LITTORINA SAXATILIS. Journal of Molluscan Studies, 71(4), 313–318. 10.1093/mollus/eyi037

Christmas, M. J., Jones, J. C., Olsson, A., Wallerman, O., Bunikis, I., Kierczak, M., Whitley, K. M., Sullivan, I., Geib, J. C., Miller-Struttmann, N. E., & Webster, M. T. (2022). A genomic and morphometric analysis of alpine bumblebees: Ongoing reductions in tongue length but no clear genetic component. Molecular Ecology, 31(4), 1111–1127. 10.1111/mec.16291

Clark, M. S., Thorne, M. A. S., Amaral, A., Vieira, F., Batista, F. M., Reis, J., & Power, D. M. (2013). Identification of molecular and physiological responses to chronic environmental challenge in an invasive species: The Pacific oyster, Crassostrea gigas. Ecology and Evolution, 3(10), 3283–3297. 10.1002/ece3.719

Collyer, M. L., & Adams, D. C. (2018). RRPP: An R package for fitting linear models to high-dimensional data using residual randomization. Methods in Ecology and Evolution, 9(7), 1772–1779. 10.1111/2041-210X.13029

Collyer, M. L., & Adams, D. C. (2024). Interrogating Random and Systematic Measurement Error in Morphometric Data. Evolutionary Biology, 51(1), 179–207. 10.1007/s11692-024-09627-6

Conde-Padín, P., Grahame, J. W., & Rolán-Alvarez, E. (2007). Detecting shape differences in species of the Littorina saxatilis complex by morphometric analysis. Journal of Molluscan Studies, 73(2), 147–154. 10.1093/mollus/eym009

Contreras-Figueroa, G., & Aragón, J. L. (2023). A Mathematical Model for Mollusc Shells Based on Parametric Surfaces and the Construction of Theoretical Morphospaces. Diversity, 15(3), 431. 10.3390/d15030431

Doyle, D., Gammell, M. P., & Nash, R. (2018). Morphometric methods for the analysis and classification of gastropods: A comparison using Littorina littorea. Journal of Molluscan Studies, 84(2), 190–197. 10.1093/mollus/eyy010

Dytham, C., Mill, P. J., Grahame, J., & O’Higgins, P. (1992). Fourier analysis as a size-free description of shape for rough periwinkle shells. Fourier Analysis as a Size-Free Description of Shape for Rough Periwinkle Shells, 127–133.

Evans, A. R., Pollock, T. I., Cleuren, S. G. C., Parker, W. M. G., Richards, H. L., Garland, K. L. S., Fitzgerald, E. M. G., Wilson, T. E., Hocking, D. P., & Adams, J. W. (2021). A universal power law for modelling the growth and form of teeth, claws, horns, thorns, beaks, and shells. BMC Biology, 19(1), 58. 10.1186/s12915-021-00990-w

Ferson, S., Rohlf, F. J., & Koehn, R. K. (1985). Measuring Shape Variation of Two-dimensional Outlines. Systematic Biology, 34(1), 59–68. 10.1093/sysbio/34.1.59

Fox, N. S., Veneracion, J. J., & Blois, J. L. (2020). Are geometric morphometric analyses replicable? Evaluating landmark measurement error and its impact on extant and fossil Microtus classification. Ecology and Evolution, 10(7), 3260–3275. 10.1002/ece3.6063

Fraley, C., Raftery, A. E., Scrucca, L., Murphy, T. B., & Fop, M. (2017). *mclust: Gaussian Mixture Modelling for Model-Based Clustering, Classification, and Density Estimation* (Version 5.4) [Computer software]. https://CRAN.R-project.org/package=mclust

Fruciano, C. (2016). Measurement error in geometric morphometrics. Development Genes and Evolution, 226(3), 139–158. 10.1007/s00427-016-0537-4

Galindo, J., & Grahame, J. W. (2014). Ecological Speciation and the Intertidal Snail Littorina saxatilis. Advances in Ecology, 2014(1), 239251. 10.1155/2014/239251

Galindo, J., Morán, P., & Rolán-Alvarez, E. (2009). Comparing geographical genetic differentiation between candidate and noncandidate loci for adaptation strengthens support for parallel ecological divergence in the marine snail Littorina saxatilis. Molecular Ecology, 18(5), 919–930. 10.1111/j.1365-294X.2008.04076.x

Garcia Castillo, D., Barton, N., Faria, R., Larsson, J., Stankowski, S., Butlin, R., Johannesson, K., & Westram, A. M. (2024). Predicting rapid adaptation in time from adaptation in space: A 30-year field experiment in marine snails. Science Advances, 10(41), eadp2102. 10.1126/sciadv.adp2102

Gefaell, J., Varela, N., & Rolán-Alvarez, E. (2020). Comparing shape along growth trajectories in two marine snail ecotypes of Littorina saxatilis: A test of evolution by paedomorphosis. Journal of Molluscan Studies, 86(4), 382–388. 10.1093/mollus/eyaa020

Gower, J. C. (1975). Generalized procrustes analysis. Psychometrika, 40(1), 33–51.

Grahame, J., & Mill, P. J. (1989). Shell Shape Variation in Littorina Saxatilis and L. Arcana: A Case of Character Displacement? Journal of the Marine Biological Association of the United Kingdom, 69(4), 837–855. 10.1017/S0025315400032203

Haines, A. J., & Crampton, J. S. (2000). Improvements To The Method Of Fourier Shape Analysis As Applied In Morphometric Studies. Palaeontology, 43(4), 765–783. 10.1111/1475-4983.00148

Hollander, J., Collyer, M. L., Adams, D. C., & Johannesson, K. (2006). Phenotypic plasticity in two marine snails: Constraints superseding life history. Journal of Evolutionary Biology, 19(6), 1861–1872. 10.1111/j.1420-9101.2006.01171.x

Jackson, H. J., Larsson, J., & Davison, A. (2021). Quantitative measures and 3D shell models reveal interactions between bands and their position on growing snail shells. Ecology and Evolution, 11(11), 6634–6648. 10.1002/ece3.7517

Johannesson, B. (1986). Shell morphology of Littorina saxatilis Olivi: The relative importance of physical factors and predation. Journal of Experimental Marine Biology and Ecology, 102(2), 183–195. 10.1016/0022-0981(86)90175-9

Johannesson, K. (2016). What can be learnt from a snail? Evolutionary Applications, 9(1), 153–165. 10.1111/eva.12277

Johannesson, K., Faria, R., Moan, A. L., Rafajlović, M., Westram, A. M., Butlin, R. K., & Stankowski, S. (2024). Diverse pathways to speciation revealed by marine snails. Trends in Genetics, 40(4), 337–351. 10.1016/j.tig.2024.01.002

Johannesson, K., Panova, M., Kemppainen, P., André, C., Rolán-Alvarez, E., & Butlin, R. K. (2010). Repeated evolution of reproductive isolation in a marine snail: Unveiling mechanisms of speciation. Philosophical Transactions of the Royal Society B: Biological Sciences, 365(1547), 1735–1747. 10.1098/rstb.2009.0256

Johnson, A. B., Fogel, N. S., & Lambert, J. D. (2019). Growth and morphogenesis of the gastropod shell. Proceedings of the National Academy of Sciences of the United States of America, 116(14), 6878–6883. 10.1073/pnas.1816089116

Kess, T., & Boulding, E. G. (2019). Genome-wide association analyses reveal polygenic genomic architecture underlying divergent shell morphology in Spanish Littorina saxatilis ecotypes. Ecology and Evolution, 9(17), 9427–9441. 10.1002/ece3.5378

Koch, E. L., Morales, H. E., Larsson, J., Westram, A. M., Faria, R., Lemmon, A. R., Lemmon, E. M., Johannesson, K., & Butlin, R. K. (2022). Genetic variation for adaptive traits is associated with polymorphic inversions in Littorina saxatilis. Evolution Letters, 5(3), 196–213. 10.1002/evl3.227

Kocot, K. M., Aguilera, F., McDougall, C., Jackson, D. J., & Degnan, B. M. (2016). Sea shell diversity and rapidly evolving secretomes: Insights into the evolution of biomineralization. Frontiers in Zoology, 13(1), 23. 10.1186/s12983-016-0155-z

Laffont, R., Firmat, C., Alibert, P., David, B., Montuire, S., & Saucède, T. (2011). Biodiversity and evolution in the light of morphometrics: From patterns to processes. Comptes Rendus Palevol, La Notion d’espèce En Paléontologie : Ontogenèse, Variabilité, Évolution, 10(2), 133–142. 10.1016/j.crpv.2010.10.004

Larsson, J., Westram, A. M., Bengmark, S., Lundh, T., & Butlin, R. K. (2020). A developmentally descriptive method for quantifying shape in gastropod shells. Journal of The Royal Society Interface, 17(163), 20190721. 10.1098/rsif.2019.0721

Le Moan, A., Stankowski, S., Rafajlović, M., Ortega-Martinez, O., Faria, R., Butlin, R. K., & Johannesson, K. (2024). Coupling of twelve putative chromosomal inversions maintains a strong barrier to gene flow between snail ecotypes. Evolution Letters, 8(4), 575–586. 10.1093/evlett/qrae014

Le Pennec, G., Butlin, R. K., Jonsson, P. R., Larsson, A. I., Lindborg, J., Bergström, E., Westram, A. M., & Johannesson, K. (2017). Adaptation to dislodgement risk on wave-swept rocky shores in the snail Littorina saxatilis. PLoS ONE, 12(10), e0186901. 10.1371/journal.pone.0186901

Lishchenko, F., & Jones, J. B. (2021). Application of Shape Analyses to Recording Structures of Marine Organisms for Stock Discrimination and Taxonomic Purposes. Frontiers in Marine Science, 8. 10.3389/fmars.2021.667183

Mayk, D., Peck, L. S., & Harper, E. M. (2022). Evidence for Carbonate System Mediated Shape Shift in an Intertidal Predatory Gastropod. Frontiers in Marine Science, 9. 10.3389/fmars.2022.894182

Mitteroecker, P., & Gunz, P. (2009). Advances in Geometric Morphometrics. Evolutionary Biology, 36(2), 235–247. 10.1007/s11692-009-9055-x

Morales, H. E., Faria, R., Johannesson, K., Larsson, T., Panova, M., Westram, A. M., & Butlin, R. K. (2019). Genomic architecture of parallel ecological divergence: Beyond a single environmental contrast. Science Advances, 5(12), eaav9963. 10.1126/sciadv.aav9963

Mosleh, S., Choi, G. P. T., Musser, G. M., James, H. F., Abzhanov, A., & Mahadevan, L. (2023). Beak morphometry and morphogenesis across avian radiations. Proceedings of the Royal Society B: Biological Sciences, 290(2007), 20230420. 10.1098/rspb.2023.0420

Olsen, A. M., & Westneat, M. W. (2015). StereoMorph: An R package for the collection of 3D landmarks and curves using a stereo camera set-up. Methods in Ecology and Evolution, 6(3), 351–356. 10.1111/2041-210X.12326

Pilakouta, N., Humble, J. L., Hill, I. D. C., Arthur, J., Costa, A. P. B., Smith, B. A., Kristjánsson, B. K., Skúlason, S., Killen, S. S., Lindström, J., Metcalfe, N. B., & Parsons, K. J. (2023). Testing the predictability of morphological evolution in contrasting thermal environments. Evolution, 77(1), 239–253. 10.1093/evolut/qpac018

Quenu, M., Trewick, S. A., Brescia, F., & Morgan-Richards, M. (2020). Geometric morphometrics and machine learning challenge currently accepted species limits of the land snail Placostylus (Pulmonata: Bothriembryontidae) on the Isle of Pines, New Caledonia. Journal of Molluscan Studies, 86(1), 35–41. 10.1093/mollus/eyz031

Ramírez-Portilla, C., Bieger, I. M., Belleman, R. G., Wilke, T., Flot, J.-F., Baird, A. H., Harii, S., Sinniger, F., & Kaandorp, J. A. (2022). Quantitative three-dimensional morphological analysis supports species discrimination in complex-shaped and taxonomically challenging corals. Frontiers in Marine Science, 9. 10.3389/fmars.2022.955582

Ravinet, M., Westram, A., Johannesson, K., Butlin, R., André, C., & Panova, M. (2016). Shared and nonshared genomic divergence in parallel ecotypes of Littorina saxatilis at a local scale. Molecular Ecology, 25(1), 287–305. 10.1111/mec.13332

Renaud, S., Michaux, J., Schmidt, D. N., Aguilar, J.-P., Mein, P., & Auffray, J.-C. (2005). Morphological evolution, ecological diversification and climate change in rodents. Proceedings of the Royal Society B: Biological Sciences, 272(1563), 609–617. 10.1098/rspb.2004.2992

Robinson, C. V., & Visona-Kelly, B. C. (2025). A geometric morphometric approach for detecting different reproductive stages of a free-ranging killer whale Orcinus orca population. Scientific Reports, 15(1), 3239. 10.1038/s41598-025-86793-3

Rohlf, F. (1998). Geometric Morphometrics and Phylogeny. Systematic Biology, 47, 147–158; discussion 159. 10.1080/106351598261094

Rolán-Alvarez, E. (2007). Sympatric speciation as a by-product of ecological adaptation in the Galician Littorina saxatilis hybrid zone. Journal of Molluscan Studies, 73(1), 1–10. 10.1093/mollus/eyl023

Scrucca, L., & Raftery, A. E. (2018). clustvarsel: A package implementing variable selection for Gaussian model-based clustering in R. Journal of Statistical Software, 84, 1–28.

Serra, F., Balseiro, D., Monnet, C., Randolfe, E., Bignon, A., Rustán, J. J., Bault, V., Muñoz, D. F., Vaccari, N. E., Martinetto, M., Crônier, C., & Waisfeld, B. G. (2023). A dynamic and collaborative database for morphogeometric information of trilobites. Scientific Data, 10, 841. 10.1038/s41597-023-02724-9

Steinley, D. (2004). Properties of the hubert-arable adjusted rand index. Psychological Methods, 9(3), 386.

Stoffel, M. A., Nakagawa, S., & Schielzeth, H. (2017). rptR: Repeatability estimation and variance decomposition by generalized linear mixed-effects models. Methods in Ecology and Evolution, 8(11), 1639–1644. 10.1111/2041-210X.12797

Tirado, T., Saura, M., Rolán-Alvarez, E., & Quesada, H. (2016). Historical Biogeography of the Marine Snail Littorina saxatilis Inferred from Haplotype and Shell Morphology Evolution in NW Spain. PLOS ONE, 11(8), e0161287. 10.1371/journal.pone.0161287

Westbury, M. V., Brown, S. C., Lorenzen, J., O’Neill, S., Scott, M. B., McCuaig, J., Cheung, C., Armstrong, E., Valdes, P. J., Samaniego Castruita, J. A., Cabrera, A. A., Blom, S. K., Dietz, R., Sonne, C., Louis, M., Galatius, A., Fordham, D. A., Ribeiro, S., Szpak, P., & Lorenzen, E. D. (2023). Impact of Holocene environmental change on the evolutionary ecology of an Arctic top predator. Science Advances, 9(45), eadf3326. 10.1126/sciadv.adf3326

Westram, A. M., Rafajlović, M., Chaube, P., Faria, R., Larsson, T., Panova, M., Ravinet, M., Blomberg, A., Mehlig, B., Johannesson, K., & Butlin, R. (2018). Clines on the seashore: The genomic architecture underlying rapid divergence in the face of gene flow. Evolution Letters, 2(4), 297–309. 10.1002/evl3.74

Yan, M., Chen, X., Xue, J., Liu, H., Jiang, T., & Yang, J. (2024). Copper Stress Causes Shell Morphology Changes in Early Juvenile Anodonta woodiana Based on Geometric–Morphometric Analysis. Bulletin of Environmental Contamination and Toxicology, 112(2), 28. 10.1007/s00128-024-03855-4

