## Supplementary Figure 1 for "Benchmarking Geometric Morphometric Methods: A Performance Evaluation for Gastropod Shell Shape Analyses"

- **L1:** Apex of the shell.
- **L2:** Suture point at the upper-right junction of the body whorl and penultimate whorl.
- **L3:** Suture point at the upper-left junction of the body whorl and penultimate whorl.
- **L4:** Suture point at the lower-left junction of the body whorl and penultimate whorl.
- **L5:** Terminal point of the body whorl suture.
- **L6:** Apex of the inner curve between the shell body and the aperture.
- **L7:** Lowest point of the shell's ventral margin.
- **Line 1:** A reference line defined between L1 (apex) and L6 (inner curve apex).
- **Comb 1:** Comprised 15 branches oriented parallel to Line 1. The top branch was placed at L4, and the bottom branch was placed at L7 (lowest shell point).
- **Comb 2:** Comprised 7 branches oriented parallel to Line 1. The top branch was placed at L3, and the bottom branch was placed at L6.
- Semi-landmarks were defined as the points of intersection between the branches of Comb 1 and both the inner lip and outer visible margin of the last whorl.

- Similarly, semi-landmarks were placed as the points of intersection between the branches of Comb 2 and the external surface of the shell.

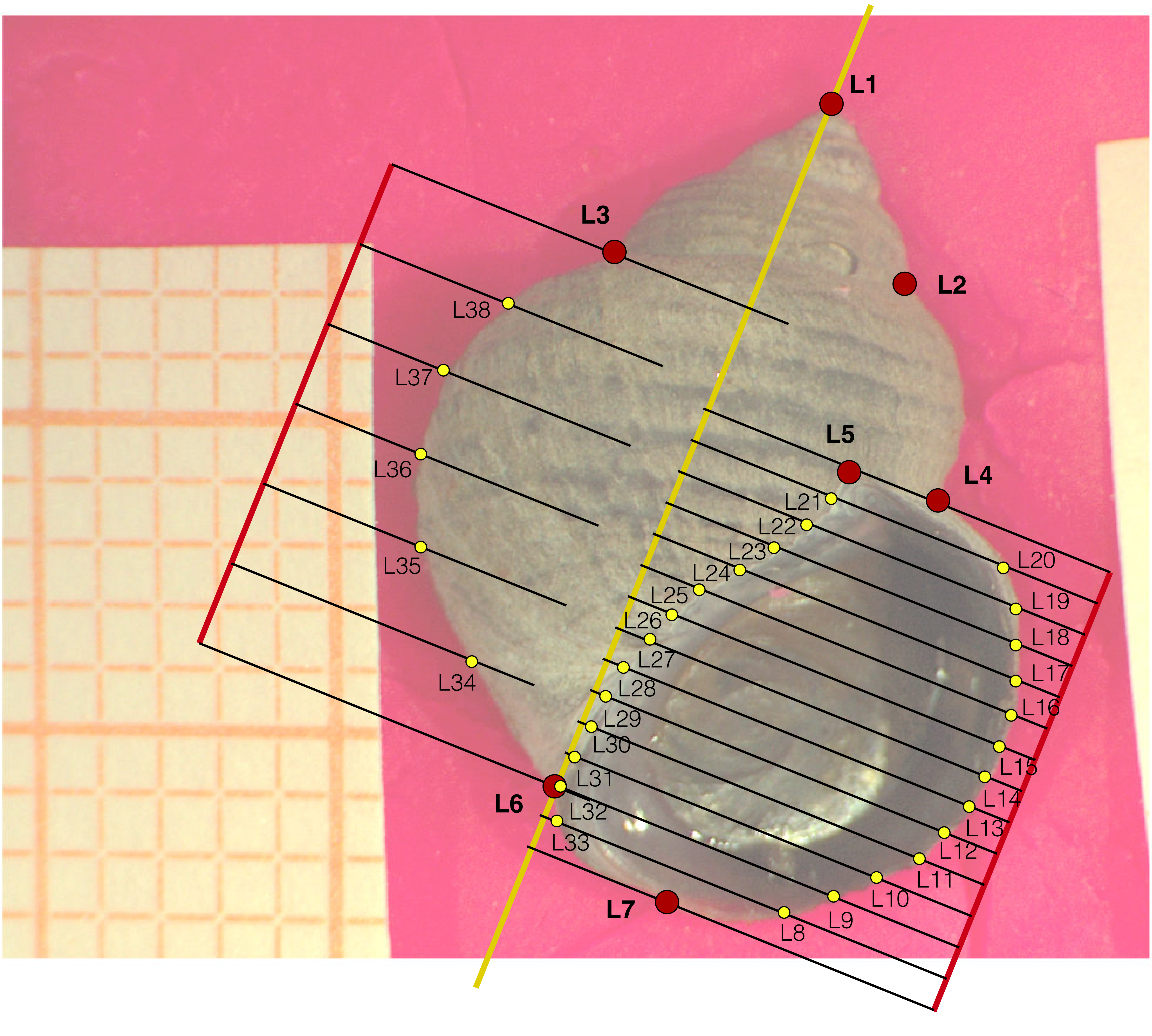


**Supplementary Figure 1:** Landmark and semi-landmark configuration for Littorina Saxatilis shell shape.
