## Supplementary figures and images for "Benchmarking Geometric Morphometric Methods: A Performance Evaluation for Gastropod Shell Shape Analyses"

### Supplementary Figure 2

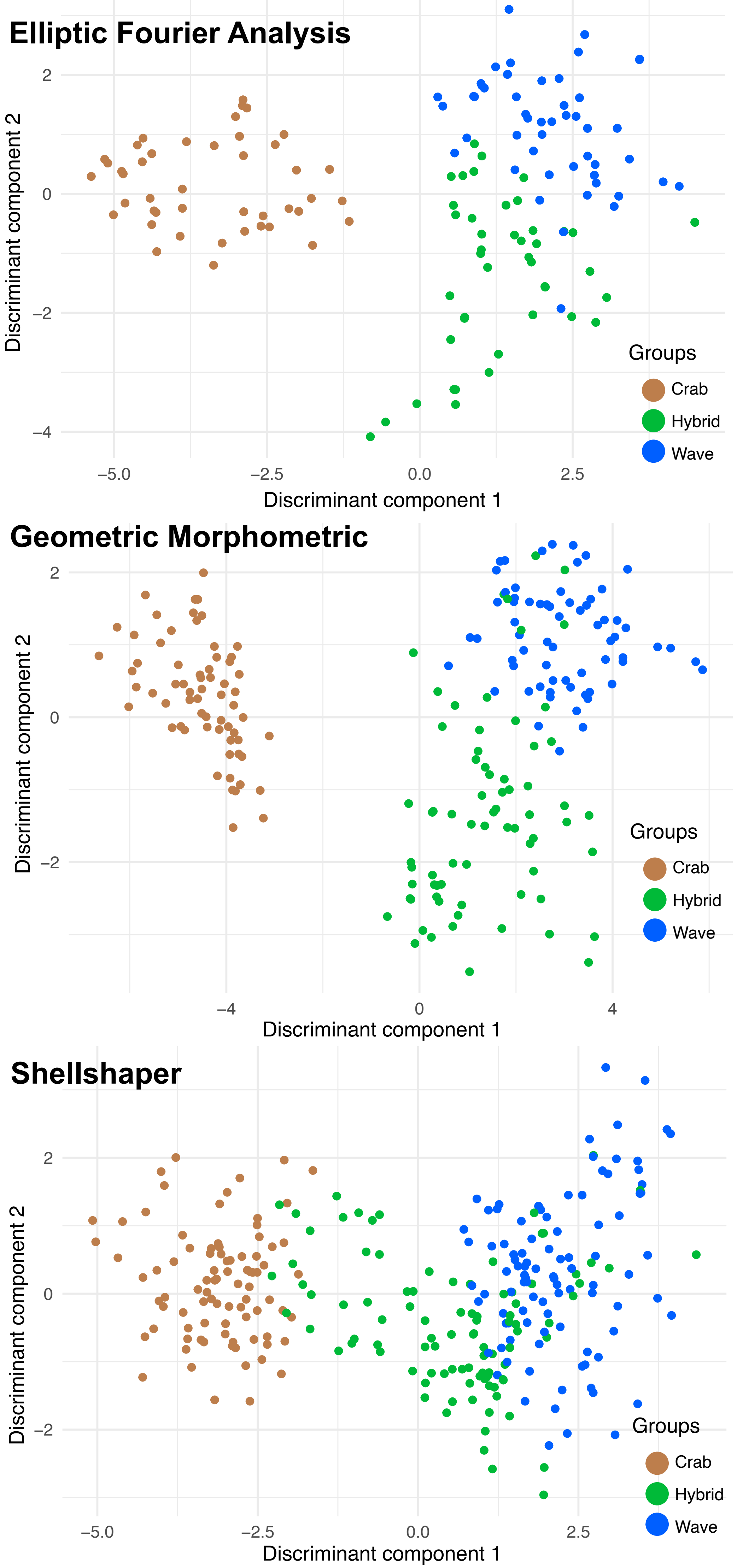


**Figure 1:** Discriminant analysis plot of EFA, GM and SS techniques
